# Emergence of Novel Lineage of Foot-and-Mouth Disease Virus Serotype Asia1 BD-18 (G-IX) in Bangladesh

**DOI:** 10.1101/604892

**Authors:** M. Rahmat Ali, A. S. M. R. U. Alam, M. Al Amin, Mohammad A. Siddique, Munawar Sultana, M. Anwar Hossain

## Abstract

In 2018, a novel circulatory foot-and-mouth disease virus serotype Asia1 BD-18 (G-IX) lineage containing a unique mutation has emerged in Bangladesh. This emergence may be following the evolutionary roadmap of previously reported lineage. Inappropriate vaccination and inefficient outbreak surveillance possibly contributed to the current episode of emergence.

## Text

Foot-and-Mouth Disease is an endemic animal disease of transboundary importance in Bangladesh (*1, 2*). This economically devastating, contagious disease is caused by FMD virus (FMDV; *Picornaviridae* family, *Aphthovirus* genus) which exists as seven serotypes (O, A, C, Asia1, SAT 1-3), each divided into topotypes, lineages, sublineages and strains (*3*). Among them, three serotypes (O, A and Asia1) have been circulating in Asia and serotype Asia1 is the least prevalent (*1, 3, 4*).

Serotype Asia1 is unique to Asia, and was first detected in India in 1951-52 and in Pakistan in 1954 (*3-5*). Previously this serotype was classified into lineages (*6*), while Valarcher et al. (2009) divided these FMDVs into groups based on 5% nucleotide differences (*4*). FAO World Reference Laboratory for FMD (WRLFMD) has currently established a standard reference list for the nomenclature of FMDV serotypes (*7*). According to that, the topotype for Asia1 serotypes is ASIA and the lineages are G-I to G-VIII (*4, 5, 8*). Interestingly, Valarcher et al. (2009) designated previous lineage D as G-III (*4*) and Subramaniam et al. (2013) designated stains circulating after 2005 under previous lineage C as G-VIII (*8*), while both lineages were endemic in Indian subcontinent (*6*). In recent years (2016-17), Sindh-08 (G-VII) is the only lineage of serotype Asia1 that is in continuous circulation in its endemic region (*9*) whereas outbreaks with other lineage G-VIII were reported sporadically (*10, 11*).

The last report of Asia1 from Bangladesh was in 2013 (*11, 12*), while all of the previous reported isolates from Bangladesh were within lineage G-VIII (*10*). After 2013, no Asia1 strains were reported from Bangladesh and India (*11, 13*). Here we report the emergence of a novel lineage of FMDV serotype Asia1 BD-18 (G-IX) in Bangladesh in 2018.

## The Study

On 18^th^ January 2018, an outbreak of FMD occurred in the BGB (Border Guards Bangladesh) Dairy Farm, Pilkhana, Dhaka. Tongue epithelium tissue samples were collected from FMD-suspected cattle, and subsequently virus isolation and generation of VP1-coding sequences were performed using standard protocols of our laboratory (*1*).

We reconstructed VP1 phylogeny by maximum likelihood method for the assorted dataset (Technical Appendix (TA) Table 1) in MEGA7 under on general time-reversible model of substitution with gamma distributed rate variation among sites with 1,000 pseudo-replicates and estimated nucleotide divergence (ND) among the established lineages of FMDV serotype Asia1. Using BEAST v2.4.5 (*14*), we accommodated the coalescent constant tree prior using a lognormal distribution in an uncorrelated relaxed molecular clock model across tree branches, and sampled out 1600 trees from 80 million iterations of Markov Chain Monte Carlo (MCMC) chain. Finally, summarized maximum clade credibility tree (MCC) was annotated and depicted in FigTree representing 95% HPD confidence intervals over the nodes to evaluate the reliability (TA).

**Table 1.** Percent nucleotide identity/divergence in VP1 encoding gene of FMDV Asia1 established groups with newly proposed group IX.

| Genetic lineages | Intergroup (G-IX) |  |
| --- | --- | --- |
|  | Nucleotide identity (%) | Nucleotide divergence (%) |
| G-I | 83.9-85.2 | 17-18.7 |
| G-II | 86.6-88.3 | 13-15.2 |
| G-III | 87-88.6 | 12.6-14.6 |
| G-IV | 84-86.6 | 15-20.4 |
| G-V | 83.6-85.6 | 16.3-18.9 |
| G-VI | 87.8-89.4 | 11.4-13.7 |
| G-VII | 82.3-84.7 | 17.5-20.6 |
| G-VIII | 90-91.8* | 8.8-10.9 |

\*The least identity/divergence has been observed with G-VIII. From the perspective of our dataset, two strains (IND227/07 & IND12/07) of G-VIII from different states of India showed the highest identity (~92%) to the novel G-IX lineage. Both of the virus strains had tracked back to an Indian origin in about 2005.

The proposed lineage BD-18 (G-IX) showed at least about >8% nucleotide divergence with all the eight established lineages of Asia1 serotype and highest sequence identity value of its closest genetic lineage G-VIII (Table 1). Moreover, the clades of G-VIII and G-IX separated in the distance tree with an ND of >5% demonstrating the presence of two separate lineages (TA Figure 2). The significant nucleotide variation of BD-18 with other lineages validated the existence of an independently evolving lineage within Bangladesh. The phylogenetic tree clearly represented that all the established lineages and novel lineage BD-18 (G-IX) formed distinct clades signifying the variation within lineages with well-supported bootstrap values of the key nodes (Figure 1, Panel A). Before the emergence of this lineage, lineage G-VIII was sporadically circulating in Bangladesh and its neighboring countries (*10, 11, 13*). Remarkably, the current vaccine strain (IND63/72) frequently used in Bangladesh is distantly related to newly emerged G-IX in the phylogenetic trees with ND of 16.5-16.7%.

**Figure 1.**
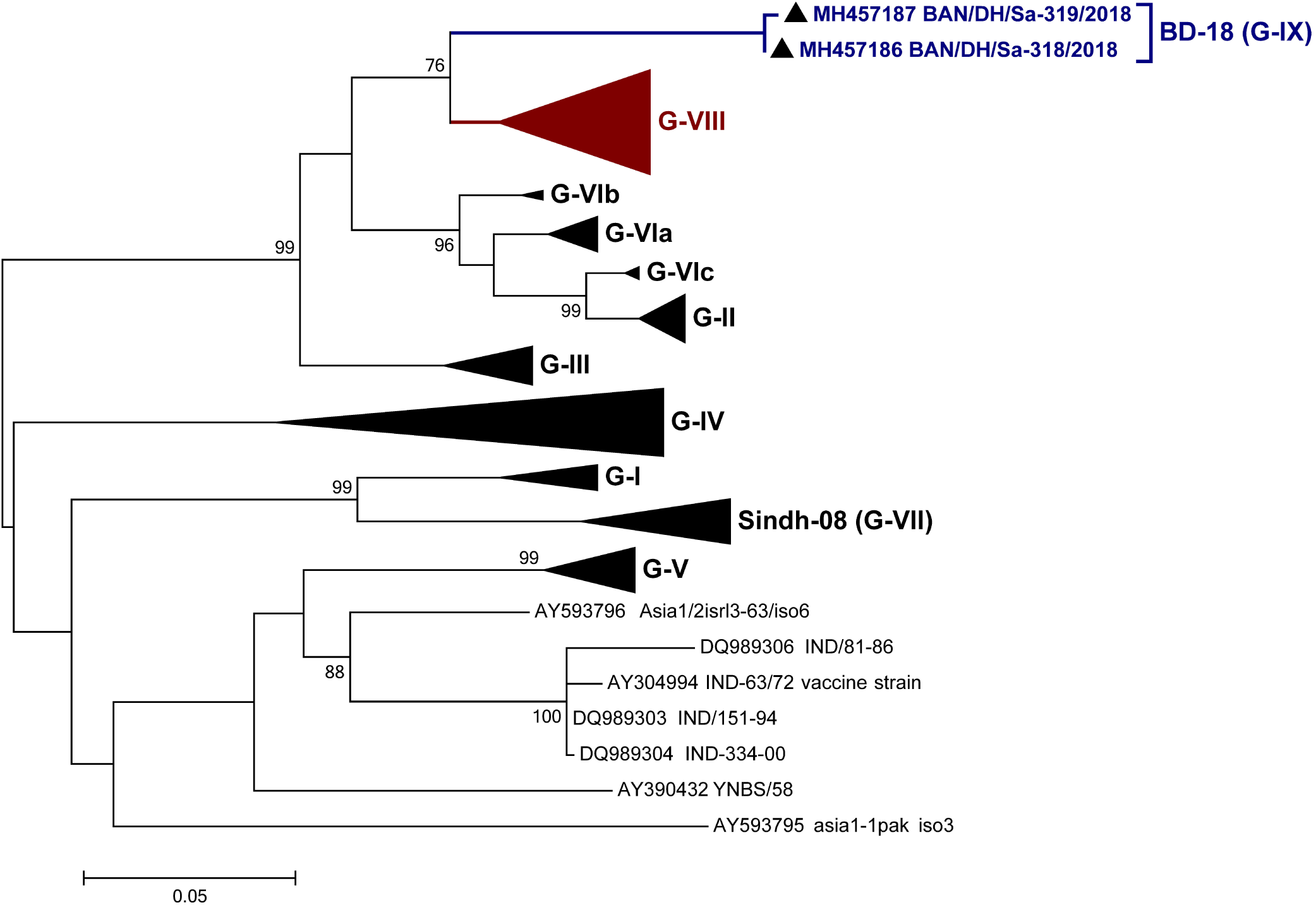
Phylogenetic analyses of VP1 coding region of serotype Asia1 covering all the lineages. A) Phylogenetic relationship among the established lineages of FMDV Asia1 serotype and novel lineage G-IX. Scale bar indicates nucleotide substitution per site. The sequences generated for this study are shown in bold letter and marked with (filled triangle) symbol. Lineage BD-18 (G-IX) formed a completely different clade seemingly originating from G-VIII, which was in circulation in Bangladesh during 2012-2013 and never found in India and/or Bangladesh after 2013. B) Maximum clade credibility (MCC) tree summary of spatio-temporal reconstruction based on VP1 sequences of FMDV serotype Asia1. The time-tree (tips corresponding to year of sampling) was generated using TreeAnnotator in BEAST2 package and annotated in FigTree with discrete location (country) based depiction. Some terminal nodes were collapsed for clarity and represented the established genetic lineages. Internal node colors reflect inferred locations for the clades, while tip and branch colors represent the sampling locations for tip branches. Diameters of internal node circles represent posterior location probability values. The sequences generated by our lab for this study were shown in navy blue color and other sequences reported from Bangladesh but not from this study are marked with dark pink color. Blue lines on the node indicate 95% high posterior density of the time of the most recent common ancestor (tMRCA). The tree clearly represents the evolutionary changes throughout the time span (1921-2018) and G-IX emerged as the last group probably from G-VIII.

**Figure 2.**
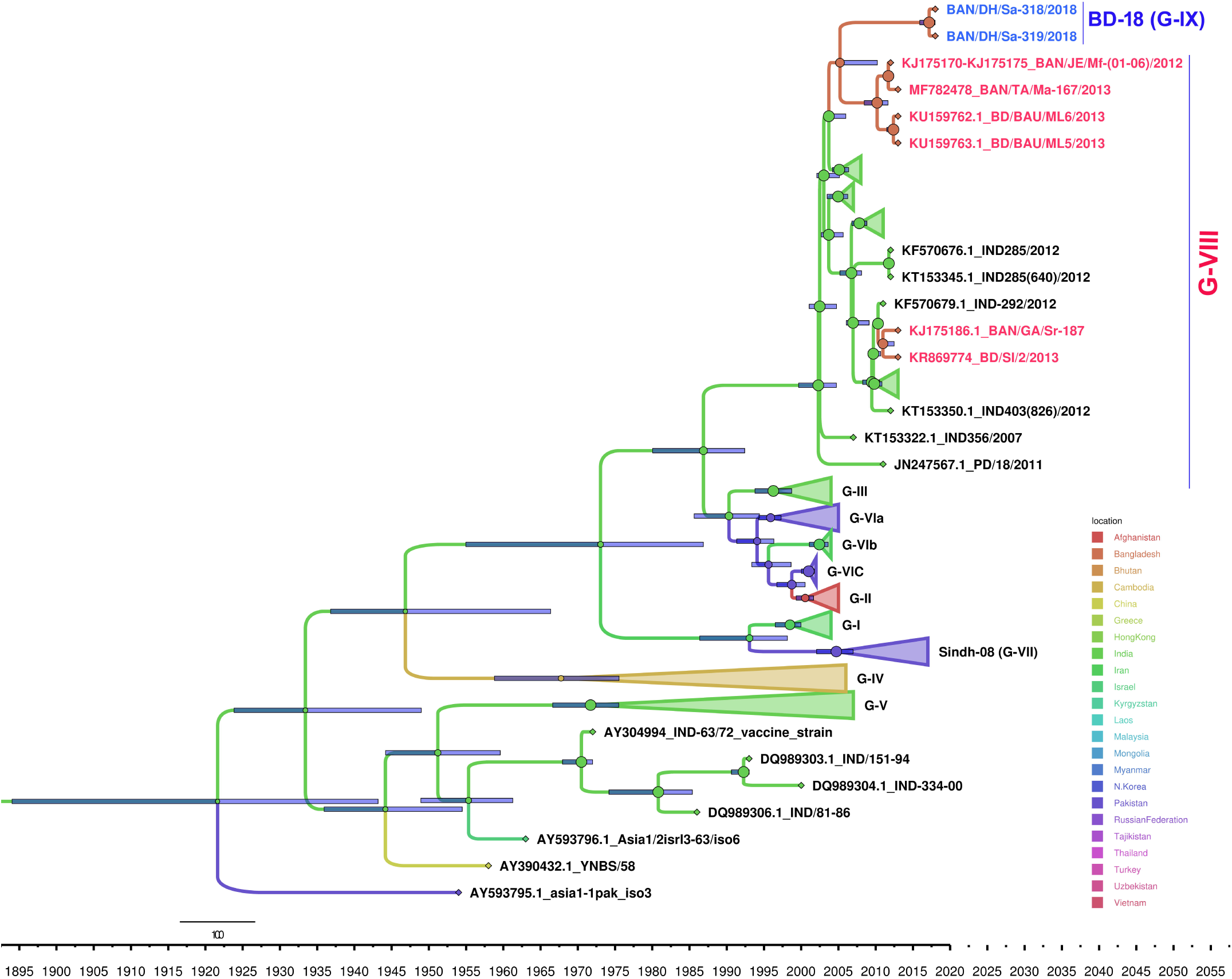
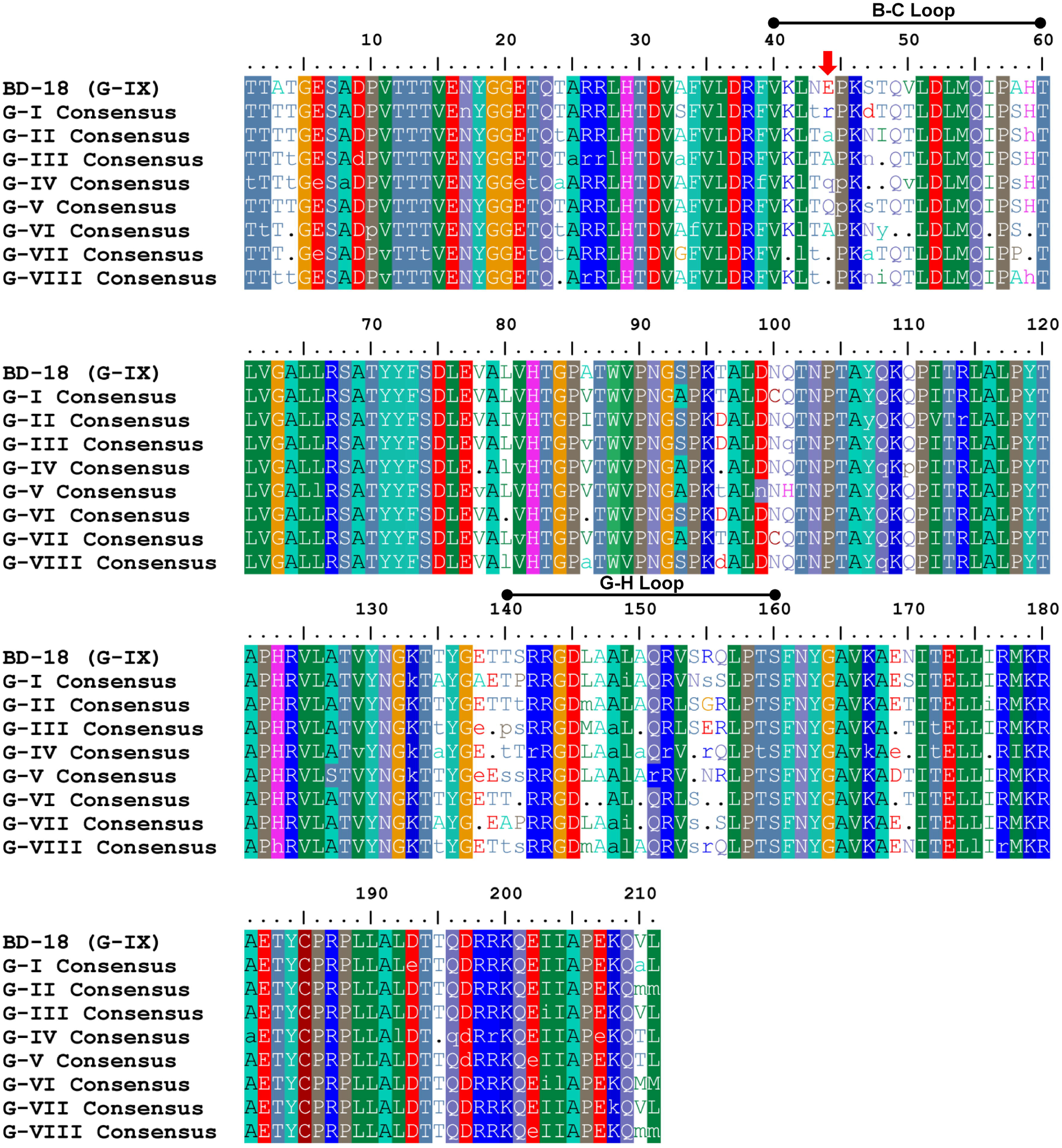
Alignment of the respective VP1 amino acid consensus of eight groups under the umbrella of FMDV Asia 1 with newly proposed group IX (contains the most recent report of FMDV Asia1 in the Indian sub-continent so far). Group specific consensus was generated based on the VP1 amino acid multiple sequence alignment of the representative isolates under the flagship of a group using MView tool (EMBL-EBI) where 80% identity was taken into consideration. Amino acids that were found conserved among all the isolates of a particular group (current dataset) were represented as uppercase letter, whereas most abundant (not conserved) amino acid call at 80% identity was denoted as a lowercase letter. In contrary, the dot present in the consensus implies the amino acid variability at those positions at 80% identity ceiling. Considering the alignment of the group specific consensuses with reference to newly proposed group IX, frequent amino acid substitution was found restricted at B-C loop (position 40-60 (*6*)) and G-H loop (position 140-160 (*6*)) of the VP1. This is noteworthy that, a unique mutation at 44 position (most probably within groove region of B-C loop) of the group IX (presence of E-Glutamic Acid, a highly charged amino acid) is evident which is indicated here as down head arrow colored in red.

From the spatio-temporal dynamics of the lineages of Asia1, we can contend that both contemporaneous and time relapsing emergence of different genetic lineages (G-I to G-VII) was noticeable through chronologically distinct evolutionary phases tracing back to most recent common ancestor (MRCA) states in five countries (TA Table 4). After 15 years since the emergence of tMRCA of G-VIII in India, BD-18 (G-IX) emerged possibly around early 2017, well before the documentation of evolving new strains, possible root of which was located in Bangladesh (Figure 1, Panel B). This lineage might have emerged through a series of key founder evolutionary events with the epidemic spreading of Indian G-VIII viruses, firstly in Bangladesh by about 2004 and then there might be intermingling of G-VIII viruses within this country in subsequent years causing the genesis of G-IX (TA Figure 3). There is also strong support behind the idea that India acted as a crucial hub (posterior probability >85%) for dissemination of viruses into Bangladesh with two independent introduction events (TA Figure 3). The sequences of combined clade of G-IX and G-VIII Bangladeshi isolates evolved at rate of 5.24×10^-3^ nucleotide/site/year, which is significantly higher, compared to an overall substitution rate of Asia1 3.824×10^-3^ nucleotide/site/year.

Lineage specific consensuses of the VP1 sequences were compared (TA) which showed frequent amino acid substitutions in B-C (10 substitutions) and G-H (8 substitutions) loops, along with heterogeneity flanking the RGD motif (Figure 2). A unique mutation at position 44 of the G-IX (Glutamic Acid, negative charged) was evident in the groove region of B-C loop, while mostly Arginine (hydrophilic, positive charged) in G-I; Alanine (hydrophobic) in G-II, III and VI; Glutamine (hydrophilic) in G-IV and V; deletion in IND63/72 vaccine strains and hyper-variability in G-VII and VIII were existent. Another interesting mutation was at position 3, where hydrophobic Alanine was present in G-IX compared to hydrophilic Threonine in other lineages. Alanine was also present at positions 58 and 86, which matches only with G-VIII.

The VP1 snapshot through reconstructing evolutionary parameters and migration pathways inferred the genetic characteristics and evolutionary history that can help in the understanding of a comprehensive picture of Asia1 lineages and evolutionary process operating in this serotype. In case of the genesis of novel lineage BD-18 (G-IX), we can mention the emergence as another key turning point for Bangladesh after the re-emergence of Asia1 serotype during 2012-2013. Expectedly, the G-VIII isolates of Bangladesh revealed close genetic relationships with Indian isolates reported since 1987 that illustrated umbrella-clustering pattern among sequences of near time-period based on both nucleotide and amino acid identity within the VP1 region (*15*). This may be due to a steady transmission of viruses through porous trans-border between these countries. From India, the gene flow caused the early landmarks of incursion of G-VIII viruses into Bangladesh, and could be an important factor to the emergence of new lineage, which was not possible to determine in this study due to lack of related sequences for recent viruses circulating in the Indian subcontinent and limitation in nature of genetic dataset.

## Conclusions

This study reports the emergence of a novel lineage BD-18 (G-IX) of serotype Asia1 in Bangladesh estimated by at least >5% nucleotide divergence, along with the re-emergence of serotype Asia1 in Bangladesh after 2013. There could be cryptic movements or circulation of viruses which cast a little doubt in the provenance of novel virus strains, however, India might be the crucial hub for independent intrusions of viruses into Bangladesh. Episode of emergence of new virus lineages makes the FMD control program complex in Bangladesh.

## Acknowledgments

This work was funded by HEQEP grant number CP-3842, provided by the University Grants Commission (UGC), Government of the People’s Republic of Bangladesh, under soft loan of World Bank.

## Author Bio

Mr. Ali is a research assistant at Microbial Genetics and Bioinformatics Laboratory, Department of Microbiology, University of Dhaka. His research interests include foot-and-mouth disease viruses and their molecular characterization.

