## Supplementary Information for "Emergence of Novel Lineage of Foot-and-Mouth Disease Virus Serotype Asia1 BD-18 (G-IX) in Bangladesh"

### Technical Appendix

#### Sample Collection:

On 24th January 2018, tongue epithelium tissue samples were collected aseptically from FMD-suspected cattle of BGB Dairy Farm, Dhaka, Bangladesh, following an outbreak commenced on 18th January 2018. The samples were collected by registered veterinary doctor with the consent of the authority and all efforts were made to minimize the sufferings of animals. The protocol for sample collection from animals was approved by the Animal Experimentation Ethical Review Committee (AEERC) under Faculty of Biological Sciences, University of Dhaka. The samples were transported to the laboratory in cold condition immediately after collection and stored at -80°C until RNA extraction or further processing.

#### RNA Extraction and cDNA Synthesis:

Homogenization of tissue sample and extraction of total RNA was performed in an automated Maxwell® 16 system (Promega, USA) using the Maxwell® 16 total RNA purification kit (Promega, USA) according to the manufacturer's instruction. Immediately after extraction, the reverse transcription of extracted RNA into complementary DNA (cDNA) was performed using the GoScript™ Reverse Transcription System (Promega, USA).

#### Polymerase Chain Reaction Amplification and Sequencing of VP1:

The entire VP1 coding region was amplified by PCR using GoTaq 2x Hot Start Colorless Master Mix (Promega, USA) with either forward primer VP1UF [5′-GTACTACRCSCAGTAC-3′] (*1*) and the reverse primer NK61 [5′-GACATGTCCTCCTGCATCTG-3′] (*2*). Afterwards, PCR amplicons were purified using the Wizard® SV Gel and PCR Clean-Up System (Promega, USA) and purified PCR products were subjected to an automated cycle sequencing reaction using BigDye® Terminator v3.1 cycle sequencing kit (Applied Biosystems®, USA) according to manufacturer's instructions with the same primers used in the PCR reaction and analysis of the data was performed in ABI Genetic Analyzer (Applied Biosystems®, USA). Afterwards, the raw sequence data were assembled using SeqMan version 7.0 (DNASTAR, Inc., Madison, WI, USA) and the assembled sequences were compared with other entries from NCBI GenBank using BLAST search to reveal the serotype level identification of the samples and submitted to NCBI GenBank.

**Assortment of Sequence Dataset for Mutational and Evolutionary Analysis:**

A total of 143 VP1 Sequences (Technical Appendix Table 1) of serotype Asia1 from twenty-three Asian countries during 1954-2017 were included in the dataset, including two VP1 sequences generated in this study. The dataset was generated based on lineage specific assortment of VP1 sequence dataset covering every geographical regions and time period, including every Asia1 VP1 sequences reported from Bangladesh and all latest sequences from NCBI GenBank. The reduction from the initial sequences was based on the screening process that was necessary to leverage the final analytical process and also for better view of the generating data.

**Phylogenetic Analysis:**

A total of 143 complete VP1 coding gene sequences from all the established lineages of FMDV Asia1 serotype were taken in the phylogenetics analysis. Both heuristic program in MEGA7 (*10*) and jModelTest (version 2.1.10) package (*11*) were used for model selection by computing likelihood score out of 88 models and best fitted model was selected using lowest Akaike Information Criterion (AIC) (*12*) and Bayesian Information Criterion (BIC) (*13*) values, which pointed out GTR (generalised time reversible) (*14*) with a discrete gamma distribution of 1.46 with 4 categories and invariant proportion of 0.42 as the best fitted model to the aligned data set. Using MEGA7, maximum likelihood tree was employed to generate the phylogenetic tree with 1000 replication of the dataset for checking the robustness of the tree topology. The topography of the tree was also checked with neighbor joining and maximum clade credibility methods.

**Calculation of Genetic Divergence:**

The nucleotide sequence identity/divergence among the established lineages containing the references sequences from published articles and our proposed novel lineage as well as within the nine lineages were calculated using MegAlign module of Lasergene core suite version 7 (DNASTAR Inc., USA). We also reconstruct a phylogenetic tree based on Kimura distance formula to show the nucleotide divergence percentage between G-IX and its closest lineage VIII (Technical Appendix Table 2). We also performed MEGA7 based genetic distance measurement between the lineages.

**Bayesian Phylogenetic Inference**

For the sequence substitution process, we employed the same model as for phylogenetic reconstruction for estimating evolutionary rates and phylogeographical analysis considering 23 countries as discrete traits over a time period of 64 years calibrated in time units (year) using a molecular clock process with dated tips. Stepping stone (SS) method (*15*) for a combination of three coalescent and two relaxed clock models (*16*) was employed to calculate the marginal log-likelihood estimation (MLE) of each combination (*17*), and model with the significant Bayes Factor (BF) (*18*) as compared to other models was selected as the best model (Technical Appendix Table 3). Using BEAST v2.4.5 (*19*), the Markov Chain Monte Carlo (MCMC) chains were run (*20*) for 80 million steps with the selected coalescent tree prior and relaxed molecular clock models, sampling over 5,000 generations. The log files generated in these analyses were visualized in Tracer v1.6 (http://tree.bio.ed.ac.uk/software/tracer) after 10% burn-in and the statistical uncertainties were summarized in the 95% highest probability density (HPD) intervals (*21*), to estimate the relevant evolutionary parameters related to this study (substitution rate and tMRCA of root node). The maximum clade credibility (MCC) trees were generated using TreeAnnotator in BEAST2 package and the final tree was visualized using FigTree version 1.4.3 (available at http://tree.bio.ed.ac.uk/software).

At first, Asia1 has the tMRCA in the year of 1921 in India, which then separated into many distinct groups, and travelling throughout India as well as trespassing to other countries, such as China, Pakistan and Israel, created G-V on September 1971 from India as MRCA country spreading to Russia, Mongolia, Vietnam, North Korea and Myanmar as well (*22*). The movement of virus changed rapidly throughout the time course and covered some other countries at the same time period, such as Taiwan, Hong Kong, Cambodia, Laos and Malaysia generating another different lineage G-IV in 1967 in Cambodia, most probably from an Indian ancestor. These two events showed that two contemporary and different viral lineages were in circulation, which had close tMRCA but different ancestor states in that time period in Asia.

Then after a long period of time another new lineage G-VI emerged in 1995 from Pakistan and interestingly, another different novel genetic lineage G-III emerged after only five months later from India spreading also to Bhutan, Myanmar and Lebanon in the following years. Later, G-VI was further classified into three distinct sublineages- a, b and c and altogether this lineage covers four countries (Pakistan, Iran, Greece and Turkey) originating from mainly Pakistan and Iran territory around 2001-2002. After passing two years, another lineage G-I appeared in 1998 from Iran and for the first time, Asia1 intruded into Afghanistan. Remarkably again after a two years lapse, another distinct Asia1 lineage G-II came into action in 2000, which had probable origin in Afghanistan causing contemporaneous outbreaks in Afghanistan, Tajikistan, Kyrgyzstan, Uzbekistan, Hong Kong, Iran and Pakistan. Group-VIII consisting of isolates of mainly India along with a few from Bangladesh and Myanmar covering a broad time-scale (2005-2017), could have an origin in India by about 2002. Two year later, Sindh-08 (G-VII) might arise in 2004 from Pakistan as MRCA state, which was reported from Pakistan, Turkey and Iraq. The most interesting fact is the time lapsing for emerging of a new lineage of Asia1 that is around two years for G-I, II, VII and VIII. Interestingly, India is the MRCA state for lineages G-II, III and VI taking as a single cluster having an ancestral year around 1990.

In case of Bangladeshi isolates, the MRCA states were in Bangladesh but there were three different tMRCAs. The isolates which were under G-VIII had tMRCAs in the year of 2010 but in two different months- March (clade containing BAN/TA/Ma-167/2013, BAN/JE/Mf-(01-06)/2012, BD/BAU/ML5/2013 and BD/BAU/ML6/2013) and December (clade containing BAN/Ga/Sr-187/2013 and BD/SI/2/2013). On the other hand, novel lineage G-IX has tMRCA in March, 2017 which most probably originate from Bangladesh. The combined clade of G-VIII isolates of Bangladesh and G-IX had tMRCA in around 2005 having Bangladesh as MRCA state but the key backward node of this whole cluster showed India as location and 2003 as year. The backward node of another clade of Bangladeshi isolates of G-VIII was also in India in the year of 2010. To recapitulate, the novel lineage G-IX has an obscure origin in Bangladesh as no related sequences of this new group has been available from other countries. There could be inter-country movement which is not covered in this analysis as there were not much sequence and respective states to have a good result. Moreover, the latest report of Asia1 Group VIII having available sequence was on 23 July, 2013 from Bangladesh. Placing of different sequences from separate states could give different results. It is estimated that the substitution rate for the Asia1 sequences used in this analysis covering all the lineage was 3.824× $10^{-3}$ (95% HPD 3.067× $10^{-3}$ – 4.614× $10^{-3}$) substitution/nt/year, which is lower than evolutionary rate of the clade consisting of Bangladeshi isolates (n=11) of combined group G-VIII and G-IX had an evolutionary rate of 5.24 ×$10^{-3}$.

Mostly, India and Pakistan being the most probable MRCA states might play a key role in spreading the viruses and these center hubs could be associated with viral movements and diversification through other countries. Underlying evolutionary insights for the genesis of novel lineage and spatial dynamics related to BD-18 might occur due to steady abstruse transmission and viral diversity due to poor surveillance of this semi-extinct virus type in recent years from Bangladesh. An overall phylodynamics of all Asia1 lineages might put the tMRCAs and MRCA states contentious in recognition, but we are successful in predicting a close estimation of our focused fact.

**Analysis of VP1 Amino Acid Variations:**

To visualize amino acid variation throughout the VP1 of FMDV Asia1, group specific consensuses of the VP1 were compared with newly proposed group IX. The amino acid was interpreted in light of the standard hereditary code after codon-based alignment with MEGA7 software (*10*). To generate group specific consensus, representative sequences of the group were aligned via MView tool (*23*), hence generated the consensus at 80% identity level. Representation of the consensus sequence output of MView tool was improvised for better understanding of the site specific amino acid variation. In consensus sequence, amino acids that were found conserved among all the isolates of a particular group were represented as uppercase letter, whereas most abundant (not conserved) amino acid call at 80% identity was denoted as a lowercase letter. In contrary, the dot present in the consensus sequence implies the amino acid variability at those positions at 80% identity ceiling. To capture the change in amino acid sequence of VP1, multiple sequence alignment was performed using BioEdit (*24*) where proposed group IX was taken as reference sequence with respect to the consensus VP1 amino acid sequences of the entire representative groups. To characterize the position (position 44) where a unique mutation was found, accessibility of the residue to surface was checked via Swiss-PdbViewer (Version 4.1.0) and further cross checked via Stride Web Interface. To check the accessibility of the residue of interest to surface, homology model of the Asia1/BAN/DH/Sa-

318/2018 VP1 protein was performed via SWISS-MODEL web server. The model built for the input was retrieved as pdb file and viewed by Swiss-PdbViewer (Version 4.1.0) and Stride Web Interface.

Frequent amino acid substitutions were found in B-C (10 substitutions) and G-H loops (8 substitutions), along with heterogeneity flanking the RGD motif. This is noteworthy that a unique mutation at 44 position of the G-IX (presence of E- Glutamic Acid, negative charged amino acid) was evident while variability was present in other groups like mostly Arginine (hydrophilic, positive charged) in G-I; Alanine (hydrophobic) in G-II, III and VI; mostly Glutamine (hydrophilic) in G-IV and V; deletion in IND63/72 vaccine strains and hyper-variability in G-VII and VIII. Based on the output of two interfaces (Swiss-PdbViewer, Stride Web Interface), the position was found to present possibly on the groove region of the B-C loop. Another interesting mutation was at position 3, where hydrophobic Alanine was present in G-IX compared to the presence of hydrophilic Threonine in other lineages.  Besides, Alanine was present at positions 58 and 86, which matches only with G-VIII (previous lineage C) because other lineages have different amino acids at these positions. Another independent change was evident at position 43 (Asparagine) which did not alter the amino acid characteristics, but was found only in IRN/25/2004 (G-VIb) apart from G-IX.

**Technical Appendix Table 1.** VP1 sequences of FMDV serotype Asia1 used in this study.

| Sample ID | Accession No. | Lineage | Country | Location | Date of Collection | Host | Reference |
|---|---|---|---|---|---|---|---|
| IRN//25/2004 | DQ121120 | G-I | Iran | Oromieh, West Azerbaijan | 08-11-04 | Cattle | (*3*) |
| AFG/1/2001 | DQ121109 | G-I | Afghanistan | Dand, Kandahar | 08-02-01 | Cattle | |
| AFG/2/2001 | FJ785226 | G-I | Afghanistan | Dand, Kandahar | 10-02-01 | Cattle | |
| AFG/4/2001 | DQ121110 | G-I | Afghanistan | Dand, Kandahar | 10-02-01 | Cattle | |
| IRN/11/2001 | FJ785243 | G-I | Iran | Shimiz Abad, Markalay | 08-07-01 | Cattle | |
| IRN/4/2001 | DQ121118 | G-I | Iran | Mohamad Abad | 06-07-01 | Cattle | |
| IRN/25/2001 | FJ785244 | G-I | Iran | N/A | 2001 | N/A | |
| IRN/63/2001 | FJ785245 | G-I | Iran | Takily, Talesh, Gilan | 17-12-01 | Cattle | |
| HKN/3/2005 | FJ785235 | G-II | Hong Kong | Sheung Shui, New Territories | 09-03-05 | Cattle | (*3*) |
| HKN/4/2005 | FJ785236 | G-II | Hong Kong | Sheung Shui, New Territories | 09-03-05 | Cattle | |
| HKN/6/2005 | FJ785238 | G-II | Hong Kong | Sheung Shui, New Territories | 10-03-05 | Cattle | |

| | | | | | | | |
|---|---|---|---|---|---|---|---|
| HKN/8/2005 | FJ785240 | G-II | Hong Kong | Sheung Shui, New Territories | 10-03-05 | Cattle | |
| TAJ/1/2003 | FJ785270 | G-II | Tajikistan | Khatlonsky region | 11-10-03 | Cattle | |
| TAJ/2/2003 | FJ785271 | G-II | Tajikistan | Vakhdatsky region | 10-08-03 | Cattle | |
| TAJ/5/2004 | FJ785275 | G-II | Tajikistan | | 2004 | N/A | |
| TAJ/6/2004 | FJ785276 | G-II | Tajikistan | | 2004 | N/A | |
| KRG/1/2004 | FJ785248 | G-II | Kyrgyzstan | | 2004 | N/A | |
| AFG/40/2003 | EF457991 | G-II | Afghanistan | Kapisa | 29-12-03 | Cattle | |
| UZB/2003 | FJ785277 | G-II | Uzbekistan | | 2003 | N/A | |
| AFG/138/2004 | EF457994 | G-II | Afghanistan | Ghazni | 14-02-04 | Cattle | |
| PAK/2/2004 | FJ785264 | G-II | Pakistan | | 2004 | N/A | |
| PAK/69/2003 | DQ121127 | G-II | Pakistan | | 09-10-03 | Cattle | |
| AFG/33/2003 | EF457990 | G-II | Afghanistan | Nangahar | 17-12-03 | Cattle | |
| IND/762/2003 | DQ101240 | G-III | India | Andhra Pradesh | 2003 | Cattle | (*4*) |
| IND/763/2003 | FJ785292 | G-III | India | Andhra Pradesh | 2003 | Cattle | (*3, 5*) |
| IND/139/2002 | DQ101242 | G-III | India | Bihar | 27-02-02 | Cattle | |
| IND/97/2003 | DQ989323 | G-III | India | | 2002 | Cattle | |
| IND/423/2001 | DQ989319 | G-III | India | | 2001 | Cattle | |
| IND/61/2002 | DQ989318 | G-III | India | | 2002 | Cattle | |
| IND/354/2001 | DQ989314 | G-III | India | | 2001 | Cattle | |
| IND/438/2001 | DQ989321 | G-III | India | | 2001 | Cattle | |
| IND/328/2004 | FJ785303 | G-III | India | West Bengal | 09-03-04 | Cattle | |
| IND/388/2004 | DQ101235 | G-III | India | Gujarat | 2004 | Cattle | |
| IND/148/2001 | DQ989317 | G-III | India | Gujarat | 20-12-00 | Buffalo | |
| BHU/34/2002 | DQ121112 | G-III | Bhutan | | 2002 | | (*6*) |
| VIT/1/2006 | FJ785284 | G-IV | Vietnam | Khanh Hoa Province | 27-10-05 | Cattle | (*3*) |
| VIT/2/2006 | FJ785286 | G-IV | Vietnam | Khanh Hoa Province | 27-10-05 | Cattle | |
| VIT/8/2006 | FJ785283 | G-IV | Vietnam | Khanh Hoa Province | 27-10-05 | Cattle | |
| VIT/15/2006 | FJ785281 | G-IV | Vietnam | Khanh Hoa Province | 18-10-05 | Cattle | |
| VIT/11/2006 | FJ785289 | G-IV | Vietnam | Lao Cai Province | 11-11-05 | Cattle | |
| BR/MYA/003/2006 | EU091344 | G-IV | China | Yunnan Province | 2006 | Cattle | |
| BR/MYA/004/2006 | EU091345 | G-IV | China | Yunnan Province | 2006 | Cattle | |
| BR/MYA/001/2006 | EU091342 | G-IV | China | Yunnan Province | 2006 | Cattle | |
| BR/MYA/002/2006 | EU091343 | G-IV | China | Yunnan Province | 2006 | Cattle | |
| CAM/5/1997 | FJ785229 | G-IV | Cambodia | Siem Reap | 20-06-97 | Cattle | (*6*) |
| CAM/9/1980 | FJ785228 | G-IV | Cambodia | Siem Reap | 27-11-80 | | |
| VIT/1/1992 | FJ785280 | G-IV | Vietnam | Bihn Ba Bay | 19-10-92 | Cattle | |
| LAO/3/1998 | EU667461 | G-IV | Laos | Vientiene Municipality | 05-01-98 | Cattle | |

| | | | | | | | |
|---|---|---|---|---|---|---|---|
| LAO/1/1996 | EU667460 | G-IV | Laos | Vientiene Municipality | 12-06-96 | | |
| MAY/9/1999 | FJ785251 | G-IV | Malaysia | | 16-04-99 | Cattle | |
| MAY/8/1997 | FJ785250 | G-IV | Malaysia | | 13-08-97 | Cattle | |
| MYA/3/2000 | FJ785254 | G-IV | Myanmar | | 12-06-00 | Cattle | |
| MYA/2/1997 | FJ785253 | G-IV | Myanmar | | 23-11-97 | Cattle | |
| MYA/1/2005 | FJ785257 | G-IV | Myanmar | | 28-07-05 | Cattle | |
| TAI/1/1998 | DQ121129 | G-IV | Thailand | | 1998 | | |
| VN/LC04/2005 | GU125646 | G-IV | Vietnam | Lao Cai Province | Oct-05 | Buffalo | |
| PAK/1/1954 | AY593795 | N/A | Pakistan | Okara, Punjab | 04-03-54 | Buffalo | (*3*) |
| IND/81/1986 | DQ989306 | N/A | India | | 1986 | Cattle | (*5*) |
| IND/151/1994 | DQ989303 | N/A | India | | 1993 | Cattle | |
| IND/334/2000 | DQ989304 | N/A | India | | 2000 | Cattle | |
| IND/63/1972 (VACCINE) | AF207521 | N/A | India | Maharashtra | 1972 | Cattle | (*3*) |
| China/YNBS/58 | AY390432 | N/A | China | | 1958 | Cattle | |
| Isr/13/1963 | AY593796 | N/A | Israel | | 1963 | N/A | |
| IND/16/1976 | FJ785242 | G-V | India | Ranipet, Vellore District, Tamil Nadu | 1976 | N/A | (*3*) |
| IND/18/1980 | DQ121116 | G-V | India | Kargudy, Tamil Nadu | 20-09-80 | Cattle | |
| IND/15/1981 | DQ121117 | G-V | India | Mandakarai, Nilgiris, Tamil Nadu | 17-02-81 | Cattle | |
| NKR/2/2007 | FJ785259 | G-V | N. Korea | Pyongyang | 2007 | Cattle | |
| BR/MYA/005/2006 | EU091346 | G-V | Myanmar | Yunnan Province | 2006 | Cattle | |
| BR/MYA/007/2006 | EU091348 | G-V | Myanmar | Yunnan Province | 2006 | Cattle | |
| CHA/WHN/06 | EU887277 | G-V | China | Sichuan Province | 2006 | Pig | |
| Hebei/CHA/1/2005 | EF187274 | G-V | China | Zhangjiakou city, Hebei province | Jun-05 | Cattle | |
| Beijing/CHA/2005 | EF185303 | G-V | China | Beijing municipality, Yanqing county | May-05 | Cattle | |
| MOG/2005 | FJ785252 | G-V | Mongolia | Dornod | Aug-05 | Cattle | |
| Amursky/RUS/2005 | DQ121401 | G-V | Russian Federation | Amursky | 2005 | Cattle | |
| Khabarovsk/RUS/2005 | FJ785267 | G-V | Russian Federation | Khabarovsk | 2005 | Cattle | |
| Prymorsky/RUS/2005 | FJ785268 | G-V | Russian Federation | Prymorsky | 2005 | Cattle | |
| China/Jiangsu/2005 | EF149009 | G-V | China | Jiangsu | 2005 | Cattle | (*6*) |
| GRE/2/2000 | DQ121113 | G-VIa | Greece | Feres, Evros | 01-07-00 | N/A | (*3*) |
| TUR/3/2000 | EU553915 | G-VIa | Turkey | Tukat/Zile, Kirikkale | 2000 | Cattle | |
| TUR/6/2000 | EU553916 | G-VIa | Turkey | Sivas | 2000 | Cattle | |

| | | | | | | | |
|---|---|---|---|---|---|---|---|
| TUR/8/1999 | DQ121130 | G-VIa | Turkey | Eleskirt, Agri | 10-01-99 | Cattle | |
| TUR/10/1999 | DQ121131 | G-VIa | Turkey | Eleskirt, Agri | 10-01-99 | Cattle | |
| Asia1/Igdir/TUR/1094/07/00 | DQ296529 | G-VIa | Turkey | | 2000 | | |
| IRN/58/1999 | DQ121122 | G-VIa | Iran | Tehran | 20-06-99 | N/A | |
| PAK/19/2005 | FJ785265 | G-VIa | Pakistan | Lahore, Punjab | 25-01-05 | Cattle | |
| PAK/2/1998 | EU553914 | G-VIa | Pakistan | Lahore, Punjab | 1998 | Cattle | |
| PAK/20/2003 | DQ121126 | G-VIa | Pakistan | Lahore, Punjab | 2003 | N/A | |
| PAK/3/1998 | FJ785261 | G-VIa | Pakistan | | 1998 | Cattle | |
| IRN/10/2004 | DQ121119 | G-VIb | Iran | Damshahr, Qom, Qom | 28-09-04 | Cattle | |
| IRN/30/2004 | FJ785246 | G-VIb | Iran | | 2004 | N/A | |
| IRN/31/2004 | DQ121121 | G-VIb | Iran | | 2004 | N/A | |
| PAK/30/2002 | DQ121124 | G-VIc | Pakistan | | 31-10-02 | Buffalo | |
| PAK/31/2002 | DQ121125 | G-VIc | Pakistan | | 06-11-02 | Buffalo | |
| PAK/33/2002 | FJ785262 | G-VIc | Pakistan | | 06-11-02 | Buffalo | |
| PAK/34/2002 | FJ785263 | G-VIc | Pakistan | | 06-11-02 | Buffalo | |
| SIN/PAK/L2811/2009 | HQ439190 | Sindh-08 (G-VII) | Pakistan | Hyderabad | 07-07-09 | Buffalo | (*7*) |
| PAK/8/2008 | KY091304 | Sindh-08 (G-VII) | Pakistan | Sindh | 24-12-08 | Water Buffalo | (*4*) |
| SIN/PAK/L2952/2009 | HQ439192 | Sindh-08 (G-VII) | Pakistan | | 2009 | N/A | (*7*) |
| SIN/PAK/L5/2008 | HQ439187 | Sindh-08 (G-VII) | Pakistan | Hyderabad | 22-12-08 | Cattle | |
| TUR/13/2013 | KM268898 | Sindh-08 (G-VII) | Turkey | Oguzeli, Gaziantep | 09-02-13 | Cattle | (*8*) |
| BAL/PAK/iso-1/2011 | JX435110 | Sindh-08 (G-VII) | Pakistan | Balochistan | 01-10-11 | Cattle | |
| NIAB/PUN/PAK/156/2016 | MF115991 | Sindh-08 (G-VII) | Pakistan | Punjab | 16-06-16 | N/A | |
| NIAB/PUN/PAK/197/2016 | MF167435 | Sindh-08 (G-VII) | Pakistan | | 18-05-16 | Camel | |
| NIAB/PUN/PAK/144/2016 | MF115987 | Sindh-08 (G-VII) | Pakistan | Punjab | 12-06-16 | N/A | |
| NIAB/PUN/PAK/192/2016 | MF140442 | Sindh-08 (G-VII) | Pakistan | | 05-05-16 | Cattle | |
| NIAB/PUN/PAK/181/2016 | MF140441 | Sindh-08 (G-VII) | Pakistan | | 04-04-16 | Cattle | |
| NIAB/PUN/PAK/203/2017 | MF115990 | Sindh-08 (G-VII) | Pakistan | Punjab | 12-04-17 | N/A | |
| NIAB/PUN/PAK/207/2017 | MF140437 | Sindh-08 (G-VII) | Pakistan | | 01-03-17 | Cattle | |
| NIAB/PUN/PAK/209/2017 | MF140444 | Sindh-08 (G-VII) | Pakistan | | 01-03-17 | Goat | |

| | | | | | | | |
|---|---|---|---|---|---|---|---|
| IND/292/2012 | KF570679 | G-VIII | India | Maharashtra | 23-12-11 | Cattle | (*4*) |
| IND/305/2012 | KF570683 | G-VIII | India | Maharashtra | 19-05-12 | Cattle | |
| IND/306/2012 | KF570684 | G-VIII | India | Tamil Nadu | 2012 | Cattle | |
| IND156(335)/2013 | KT153367 | G-VIII | India | Maharashtra | 21-05-13 | Cattle | |
| IND/285/2012 | KF570676 | G-VIII | India | Maharashtra | 25-01-12 | Cattle | |
| PD/509/2010 | JN247566 | G-VIII | India | Gujarat | 07-07-10 | Cattle | |
| PD/18/2011 | JN247567 | G-VIII | India | Tripura | 01-12-10 | Cattle | PD FMD, India |
| IND/787/2009 | KF570656 | G-VIII | India | Gujarat | 07-12-09 | Cattle | (*4*) |
| IND/356/2007 | KT153322 | G-VIII | India | Gujarat | 30-06-07 | Cattle | |
| IND/341/2008 | KT153324 | G-VIII | India | Assam | 02-03-08 | Cattle | |
| IND/12/2007 | HQ224553 | G-VIII | India | Assam | 12-01-07 | Cattle | |
| BAN/TA/Ma-167/2013 | MF782478 | G-VIII | Bangladesh | Madhupur, Tangail | 23-07-13 | Cattle | (*9*) |
| IND/95/2008 | HQ224558 | G-VIII | India | Madhya Pradesh | 24-10-07 | Cattle | (*4*) |
| BAN/Ga/Sr-187/2013 | KJ175186 | G-VIII | Bangladesh | Sreepur, Gazipur | 2013 | Cattle | |
| BAN/JE/Mf-(01-06)/2012 | KJ175170-5 | G-VIII | Bangladesh | Jessore | 2012 | Cattle | |
| BD/BAU/ML5/2013 | KU159763 | G-VIII | Bangladesh | Savar, Dhaka | 09-03-13 | Cattle | |
| BD/SI/2/2013 | KR869774 | G-VIII | Bangladesh | Ghatail, Tangail | 18-07-13 | Cattle | |
| BD/BAU/ML6/2013 | KU159762 | G-VIII | Bangladesh | Bera, Pabna | 17-01-13 | Cattle | |
| IND400(822)/2012 | KT153349 | G-VIII | India | Odisha | 2012 | Cattle | (*4*, *5*) |
| IND/175/2007 | KT153321 | G-VIII | India | West Bengal | 15-03-07 | Cattle | |
| IND/96/2008 | HQ224559 | G-VIII | India | Madhya Pradesh | 24-10--07 | Cattle | |
| IND148(331)/2011 | KT153333 | G-VIII | India | Gujarat | 05-08-11 | Cattle | |
| IND148(330)/2011 | KT153332 | G-VIII | India | Gujarat | 05-08-11 | Cattle | |
| IND68(128)/2012 | KT153341 | G-VIII | India | Gujarat | 28-12-11 | Cattle | |
| IND65(140)/2010 | KT153326 | G-VIII | India | Gujarat | 02-02-10 | Cattle | |
| IND403(826)/2012 | KT153350 | G-VIII | India | Karnataka | 18-08-12 | Cattle | |
| IND162(331)/2012 | KT153343 | G-VIII | India | Karnataka | 31-03-12 | Cattle | |
| IND288(643)/2012 | KT153346 | G-VIII | India | Maharashtra | 14-02-12 | Cattle | |
| IND15(24)/2012 | KT153363 | G-VIII | India | Gujarat | 18-12-11 | Cattle | |
| IND119(223)/2012 | KT153342 | G-VIII | India | Karnataka | 19-01-12 | Cattle | |
| IND/227/2007 | HQ224555 | G-VIII | India | Uttar Pradesh | 20-09-07 | Cattle | (*4*) |
| IND/205/2010 | KF570658 | G-VIII | India | Gujarat | 27-07-10 | Cattle | |
| BAN/DH/Sa-318/2018 | MH457186 | BD-18 (G-IX) | Bangladesh | Dhaka | 24-01-18 | Cattle | This paper |
| BAN/DH/Sa-319/2018 | MH457187 | BD-18 (G-IX) | Bangladesh | Dhaka | 24-01-18 | Cattle | This paper |

**Technical Appendix Table 2:** Genetic distances among the established and proposed lineage BD-08 (G-IX).

| **Genetic lineage** | **Pairwise matrix of genetic distances** | | | | | | | |
|---|---|---|---|---|---|---|---|---|
| | G-I | G-II | G-III | G-IV | G-V | G-VI | G-VII | G-VIII |
| G-II | 0.1803 | | | | | | | |
| G-III | 0.1613 | 0.0989 | | | | | | |
| G-IV | 0.1359 | 0.1521 | 0.1274 | | | | | |
| G-V | 0.1485 | 0.1771 | 0.1397 | 0.1416 | | | | |
| G-VI | 0.1666 | 0.0345* | 0.0681 | 0.1325 | 0.1547 | | | |
| G-VII | 0.0820 | 0.1724 | 0.1383 | 0.1401 | 0.1476 | 0.1571 | | |
| G-VIII | 0.1642 | 0.0828 | 0.0872 | 0.1239 | 0.1464 | 0.0625 | 0.1465 | |
| G-IX | 0.1746 | 0.1366 | 0.1234 | 0.1323 | 0.1662 | 0.1099 | 0.1716 | 0.0786# |

*The least nucleotide variation was found between two previous established genetic lineages G-III and G-VI.

#Lineage G-IX and G-VIII has genetic distance that is more ~56% variation than the lowest divergence among established lineages.

**Technical Appendix Table 3:** Model selection using stepping-stone sampling (SS) to determine the tree prior (coalescent) model and clock for serotype Asia1.

| **Clock** | **Prior tree model** | **Marginal likelihood estimator** | **Rank** | ***ln*(BF)** |
|---|---|---|---|---|
| Relaxed clock log normal | Coalescent constant population | -10178.5453 | 1 | - |
| Relaxed clock log normal | Coalescent exponential population | -10704.0575 | 2 | -6.59 |
| Relaxed clock log normal | Coalescent bayesian population | -11165.1206 | 5 | -7.59 |
| Relaxed clock exponential | Coalescent constant population | -10995.8394 | 4 | -7.40 |
| Relaxed clock exponential | Coalescent exponential population | -11335.3139 | 6 | -7.75 |
| Relaxed clock exponential | Coalescent bayesian population | -10777.1003 | 3 | -7.09 |

**Technical Appendix Table 4:** Probable tMRCA and MRCA states along with countries of nine genetic lineages of FMDV Asia1 serotype.

| Genetic lineages | Countries | tMRCA | MRCA states |
|---|---|---|---|
| G-I | (AFG, IRN ) | June 1998 | Iran |
| G-II | (PAK, AFG, KRG, T AJ, UZB, HKN, IRN ) | July 2000 | Afghanistan |
| G-III | (IND, BHU, myanamar, lebanon ) | April 1996 | India |
| G-IV | (MYA, VN, CHINA, T AI, HKN,Cambodia, laos. malaysia) | November 1967 | Cambodia |
| G-V | (CHINA, RU S, MOG, VN, NKR, MYA, IND ,ISr) | September 1971 | India |
| G-VI | (PAK, IRN, GRE, TUR) | November 1995 | Pakistan |
| | Iran | June 2002 | Iran |
| | Pakistan | January 2001 | Pakistan |
| G-VII | (PAK,turkey, iraq) | September 2004 | Pakistan |
| G-VIII | (India 2005 -2013, Bangladesh, myanmar ) | March 2002 | India |
| G-IX | Bangladesh | March 2017 | Bangladesh |

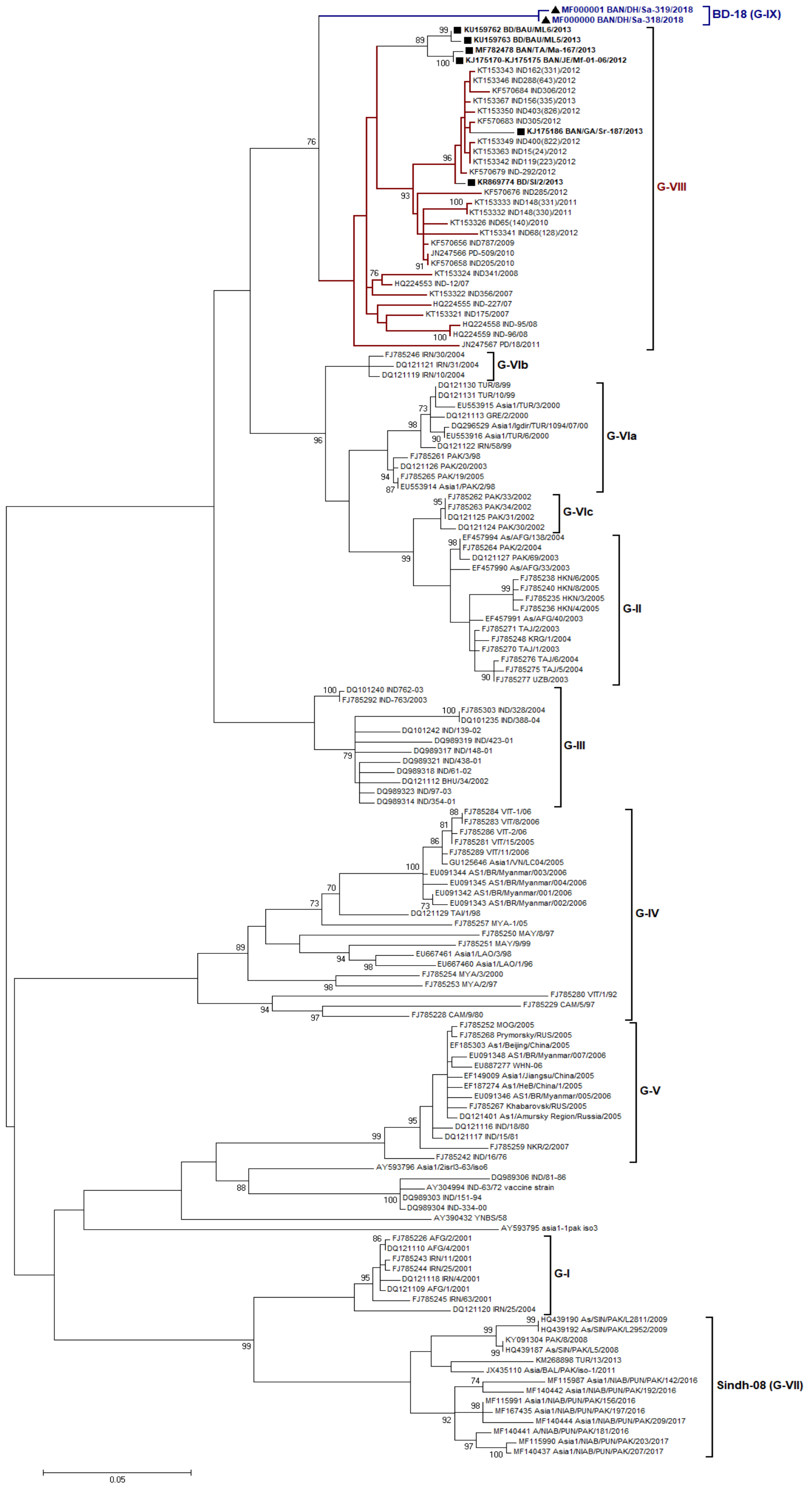


**Technical Appendix Figure 1.** Phylogenetic analyses of VP1 coding region of serotype Asia1 covering all the lineages.

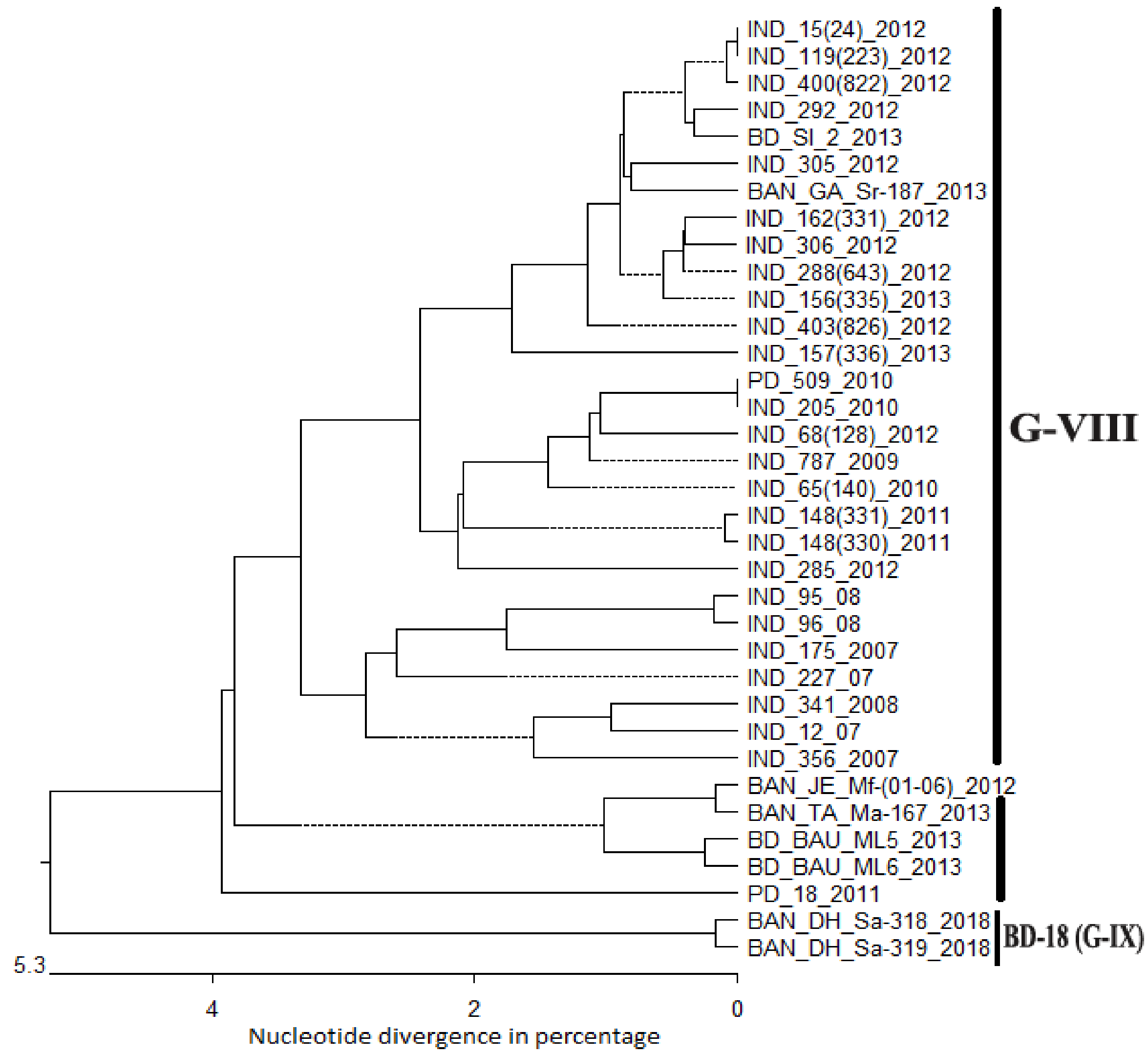


**Technical Appendix Figure 2.** The Kimura distance model based phylogenetic tree showing the nucleotide divergences (ND) among the clades of genetic lineages VIII and IX FMDV serotype Asia1. The X-axis shows the percentage nucleotide divergence (% ND). The newly proposed lineage showed >5% variation with the closest lineage G-VIII which established the group as distinct lineage.

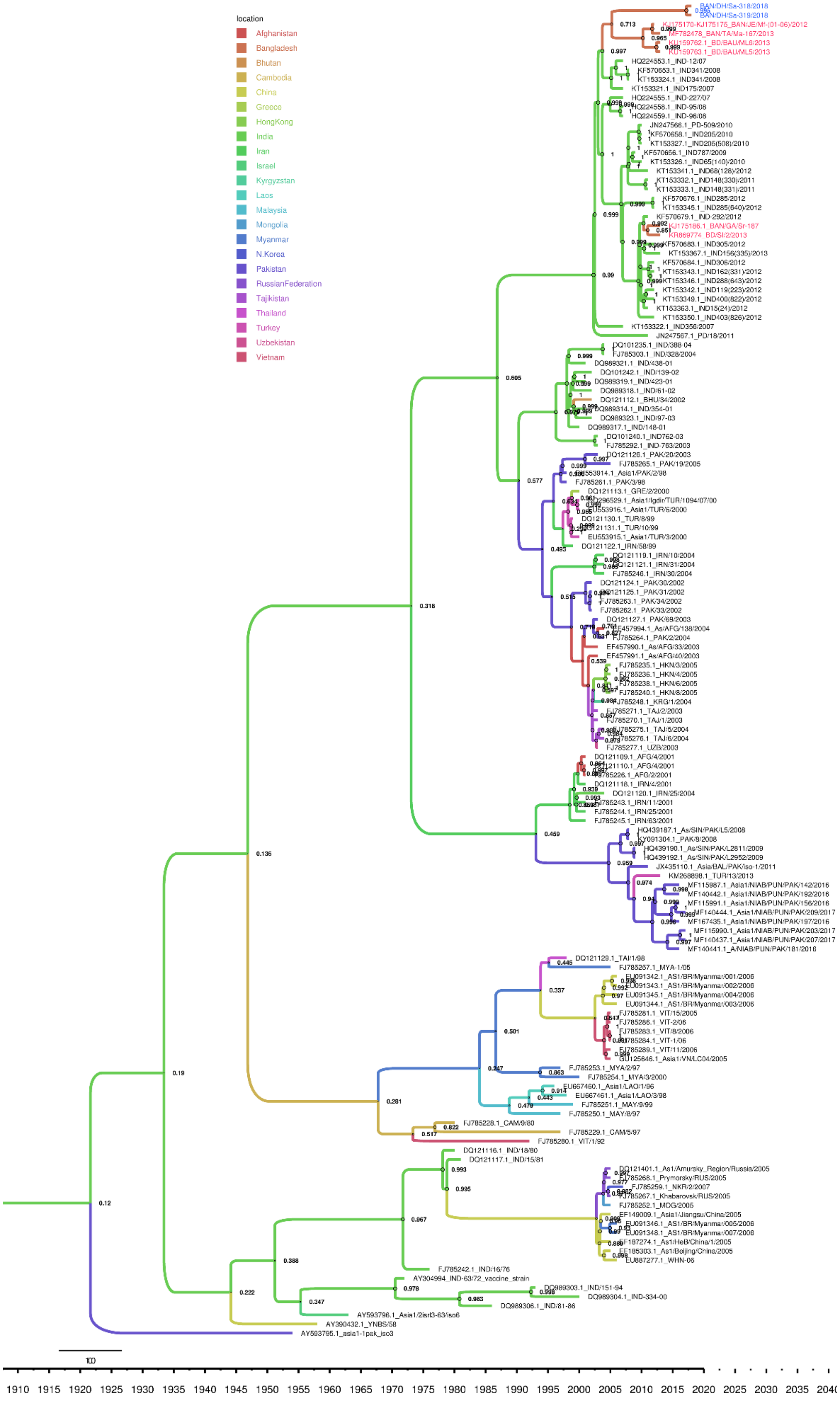
location
Afghanistan
Bangladesh
Bhutan
Cambodia
China
Greece
HongKong
India
Iran
Israel
Kyrgyzstan
Laos
Malaysia
Mongolia
Myanmar
N.Korea
Pakistan
RussianFederation
Tajikistan
Thailand
Turkey
Uzbekistan
Vietnam
100
1910 1915 1920 1925 1930 1935 1940 1945 1950 1955 1960 1965 1970 1975 1980 1985 1990 1995 2000 2005 2010 2015 2020 2025 2030 2035 2040

**Techinal appendix Figure 3.** Maximum clade credibility (MCC) tree summary of spatio-temporal reconstruction based on VP1 sequences of FMDV serotype Asia1. The time-tree (tips corresponding to year of sampling) was generated using TreeAnnotator in BEAST2 package and annotated in FigTree with discrete location (country) based depiction. Internal node colors reflect inferred locations for the clades, while tip and branch colors represent the sampling locations for tip branches. Diameters of internal node circles represent posterior location probability values with showing actual values in percentage over the nodes. The sequences generated by our lab for this study were shown in navy blue color and other sequences reported from Bangladesh but not from this study are marked with dark pink color. The tree clearly represent the evolutionary changes throughout 64 year time-scale and G-IX emerged as the last group probably from G-VIII.
